# CUT homeobox factors modulate chromatin accessibility in neurons

**DOI:** 10.64898/2026.02.10.705137

**Authors:** José Ignacio Jordá-Llorens, Jessica Valdivia, Eduardo Leyva-Díaz

## Abstract

Neuronal identity relies on coordinated transcriptional programs that include both neuron type-specific features and broadly shared pan-neuronal properties. While transcription factors that control neuronal gene expression have been defined, how these regulators interface with chromatin accessibility at neuronal loci remains poorly understood. CUT homeobox factors are broadly expressed in the *C. elegans* nervous system and provide a genetically defined system to examine transcription factor associated chromatin accessibility in neurons. Here, we establish a fluorescence-activated sorting workflow to enrich neuronal nuclei using a pan-neuronal nuclear marker and apply bulk ATAC-seq to profile chromatin accessibility in wild-type and CUT mutant animals. We identify CUT-dependent accessibility differences in neuronal nuclei, with changes concentrated at promoter-proximal regions and linked to neuronal genes, including neuronally enriched and pan-neuronal categories. Together, this work provides *in vivo* neuronal chromatin accessibility profiles and defines CUT-dependent changes at a subset of neuronal regulatory regions.

## INTRODUCTION

Neuronal identity is established and maintained by transcriptional programs that specify both neuron type-specific features and a shared set of “pan-neuronal” properties common to all neurons (Hobert et al. 2010; Hobert 2011; Deneris and Hobert 2014; Stefanakis et al. 2015). A large body of work has defined how neuron type-specific gene batteries are controlled by continuously expressed master regulators, also known as terminal selectors, that act through shared *cis*-regulatory motifs to coordinate coherent differentiation programs (Hobert 2016). In *C. elegans*, these programs have been defined with neuron type resolution and provide a tractable framework to dissect transcriptional control *in vivo* (Hobert et al. 2016). However, the regulatory logic that sustains broad pan-neuronal gene expression across diverse neuron types remains less well defined. We previously showed that CUT homeobox factors are required for robust expression of pan-neuronal genes, a shared gene expression program across the nervous system, and that CUT factors function in cooperation with master regulator transcription factors (Leyva-Diaz and Hobert 2022). Loss of CUT activity causes broad reductions in neuronal gene expression in transcriptional profiles. Together, these findings position CUT factors as broadly acting collaborators of master regulators in sustaining neuronal gene expression.

CUT homeobox genes encode DNA-binding transcription factors, including CUX and ONECUT family members (Burglin and Affolter 2016; Leyva-Diaz 2023). In *C. elegans*, six neuronal CUT genes act redundantly to support pan-neuronal gene expression, alongside master regulators (Leyva-Diaz and Hobert 2022). However, the molecular basis of this cooperation and how CUT activity interfaces with the regulatory architecture of neuronal genes remains unclear. One plausible model is that CUT activity influences the regulatory environment at neuronal loci in a way that facilitates productive engagement of additional transcription factors at shared targets, including neuron type-specific factors. Chromatin accessibility is a core feature of cell-type identity, with accessible regulatory regions strongly associated with transcription factor occupancy and gene regulatory state (Thurman et al. 2012). Consistent with a potential link between CUT family activity and chromatin state, a study in human neuronal reprogramming reported that ONECUT/CUX sequence motifs are associated with differential chromatin accessibility between fibroblasts and induced neurons, and that ONECUT overexpression is accompanied by broad accessibility changes at motif-containing regions (van der Raadt et al. 2019). These observations motivated us to test whether CUT factor activity is associated with changes in chromatin accessibility at neuronal regulatory regions in *C. elegans*.

Determining whether CUT activity is linked to chromatin accessibility calls for chromatin profiling in a neuron-enriched context, yet neuron-specific genome-wide assays in *C. elegans* remain technically challenging. The small size of *C. elegans* nuclei and the limited material available from defined cell populations have limited broader adoption of tissue-resolved chromatin profiling. While affinity-based nuclear isolation strategies can enable cell type-specific molecular profiling (Steiner et al. 2012), broader deployment of genome-wide assays has benefited from workflows that are streamlined, scalable, and compatible with rapid sample handling. Recent improvements in fluorescence-activated sorting have made it feasible to enrich fluorescently labeled nuclei at sufficient purity and yield for sequencing-based assays.

Here, we establish a workflow to enrich neuronal nuclei from *C. elegans* and use it to profile CUT-dependent chromatin accessibility *in vivo*. Using a validated pan-neuronal nuclear marker, we purified neuronal nuclei by fluorescence-activated sorting and verified nuclear integrity and neuronal enrichment. Bulk ATAC-seq profiling of sorted nuclei from wild-type and CUT mutant animals revealed widespread CUT-dependent changes in chromatin accessibility, with the majority of differentially accessible regions showing reduced accessibility in CUT mutants. Differential sites were enriched at promoter-proximal regions and were associated with genes linked to neuronal expression categories, including neuronally enriched and pan-neuronal gene sets. Together, these data provide a neuronal chromatin accessibility framework for investigating how CUT factors contribute to neuronal gene regulation.

## RESULTS

### Isolation and purification of neuronal nuclei for genomic profiling

Building on previous work showing that CUT factors are required for robust pan-neuronal gene expression (Leyva-Diaz and Hobert 2022), we profiled chromatin accessibility in neuronal nuclei to assess how CUT loss affects neuronal regulatory landscape. To define how CUT factors shape chromatin accessibility *in vivo*, we isolated neuronal nuclei from *C. elegans* and performed bulk assay for transposase-accessible chromatin (ATAC-seq) in wild-type and CUT sextuple mutant animals (lacking all six neuronal CUT homeobox genes: *ceh-44, ceh-48, ceh-38, ceh-41, ceh-21*, and *ceh-39*; hereafter, CUT mutant animals). In Leyva-Díaz and Hobert (2022), we generated and validated a CUT mutant strain carrying an INTACT-based pan-neuronal nuclear envelope reporter, which includes a mCherry-tagged nuclear envelope marker. Here, we leveraged this validated reporter to enable fluorescence-activated nuclei sorting (FANS) of neuronal nuclei (**Fig. 1A**).

**Figure 1.**
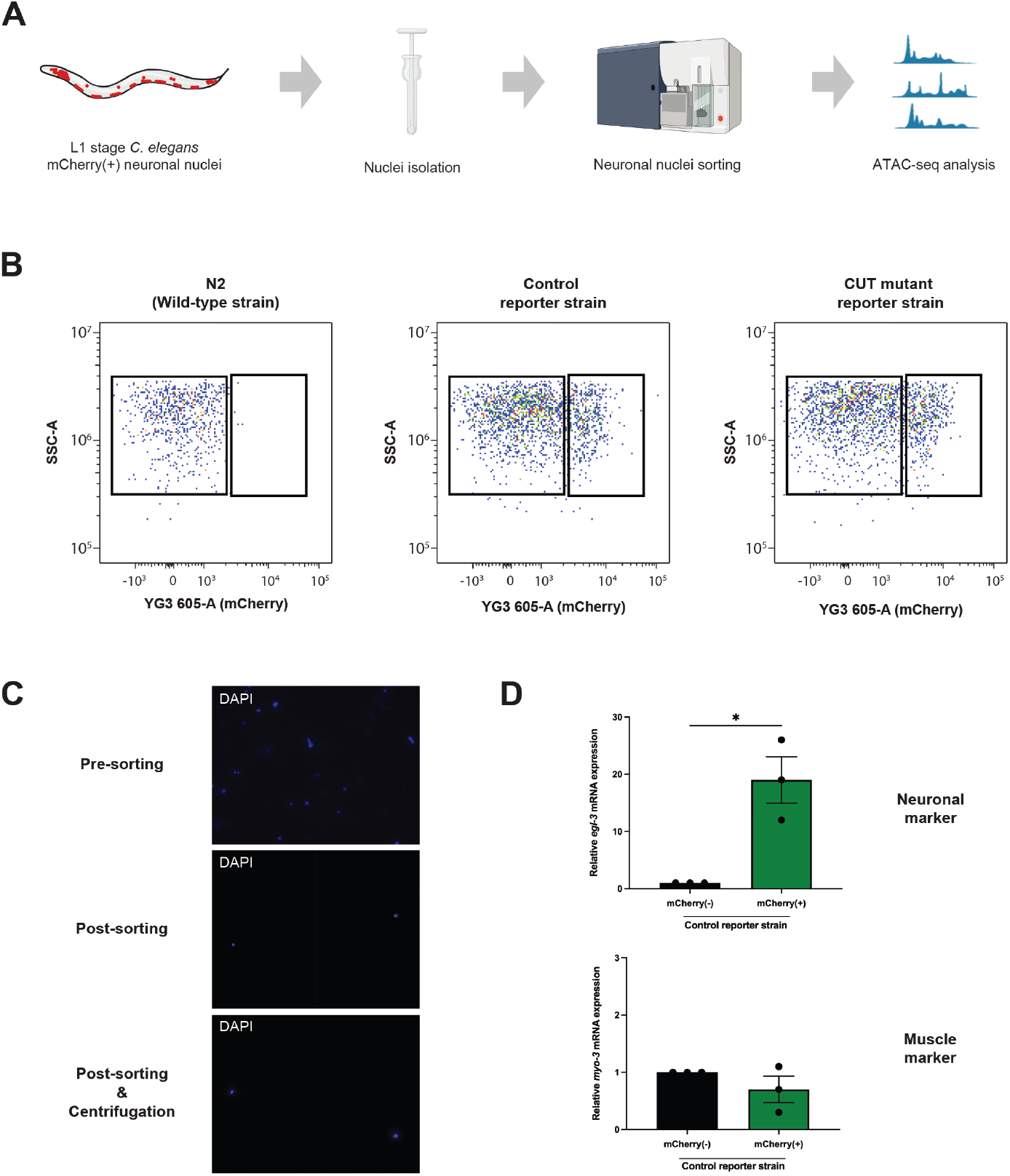
Isolation of neuronal nuclei for ATAC-seq in *C. elegans*. **(A)** Schematic overview of the workflow. L1-stage animals carrying a pan-neuronal, INTACT-based nuclear envelope reporter (mCherry-tagged) were used for nuclei extraction, fluorescence-activated sorting, and downstream ATAC-seq. **(B)** Representative flow cytometry plots (SSC-A vs mCherry fluorescence) illustrating gating for mCherry(+) (left gates) and mCherry(-) (right gates) sorted fractions. N2 (wild-type strain, no reporter) defines background fluorescence; reporter strains (Control and CUT mutant) show a distinct mCherry-high population. mCherry(+) nuclei represented 21.6% (Control) and 25.1% (CUT mutant) of total DAPI(+) events. **(C)** Representative images of DAPI-stained nuclei preparations before sorting, immediately after sorting, and after post-sort centrifugation. **(D)** RT-qPCR validation of enrichment in sorted fractions. Relative mRNA levels (ΔΔCt) are shown for a neuronal marker (*egl-3*) and a muscle marker (*myo-3*) in mCherry(-) and mCherry(+) fractions. Dots indicate independent sorts (n = 3); bars show mean ± SEM. Unpaired t test, *p < 0.05.

To allow direct comparison with our prior neuronal transcriptomic dataset, we profiled animals at the first larval (L1) stage. Nuclei were released from whole animals, stained with DAPI, and sorted by FANS using DAPI and mCherry fluorescence. Relative to N2 wild-type animals, reporter strains displayed a distinct nuclear population with high mCherry expression, and comparable mCherry(+) fractions were recovered from Control (reporter strain without CUT mutations) and CUT mutant backgrounds (**Fig. 1B**). Across independent preparations, mCherry(+) nuclei represented ∼20-25% of total DAPI(+) events in both genotypes. We applied conservative sorting gates to prioritize specificity over yield, collecting only nuclei with strong mCherry signal to minimize contamination by non-neuronal nuclei. Inspection of DAPI-stained nuclei before sorting, immediately after sorting, and after the post-sort centrifugation step used to concentrate nuclei for the tagmentation reaction indicated predominantly intact nuclear morphology throughout the workflow, supporting preservation of nuclei integrity (**Fig. 1C**). Finally, qPCR analysis of sorted material supported neuronal enrichment, with a robust increase in a pan-neuronal marker (*egl-3*) and a tendency to lower levels of a muscle marker (*myo-3*) in the mCherry(+) fraction relative to mCherry(-) nuclei (**Fig. 1D**). Together, these results establish a FANS-based workflow that yields neuron-enriched nuclei suitable for downstream genome-wide chromatin accessibility profiling.

### Neuronal ATAC-seq reveals CUT-dependent accessibility changes

Using FANS-isolated neuronal nuclei, we profiled chromatin accessibility by bulk ATAC-seq in Control and CUT mutant animals at the L1 stage. Differential accessibility analysis identified 858 differentially accessible (DA) peaks between conditions (FDR < 0.05, |log2FC| > 1). Among these, 816 peaks showed reduced accessibility and 42 showed increased accessibility in CUT mutant neuronal nuclei relative to Control (**Fig. 2A**). To assess overall ATAC-seq signal in these datasets, we examined aggregate accessibility profiles around transcription start sites (TSSs) across all peaks. Both Control and CUT mutant libraries showed signal enrichment around TSSs, reflecting promoter-proximal accessibility (**Fig. 2B**).

**Figure 2.**
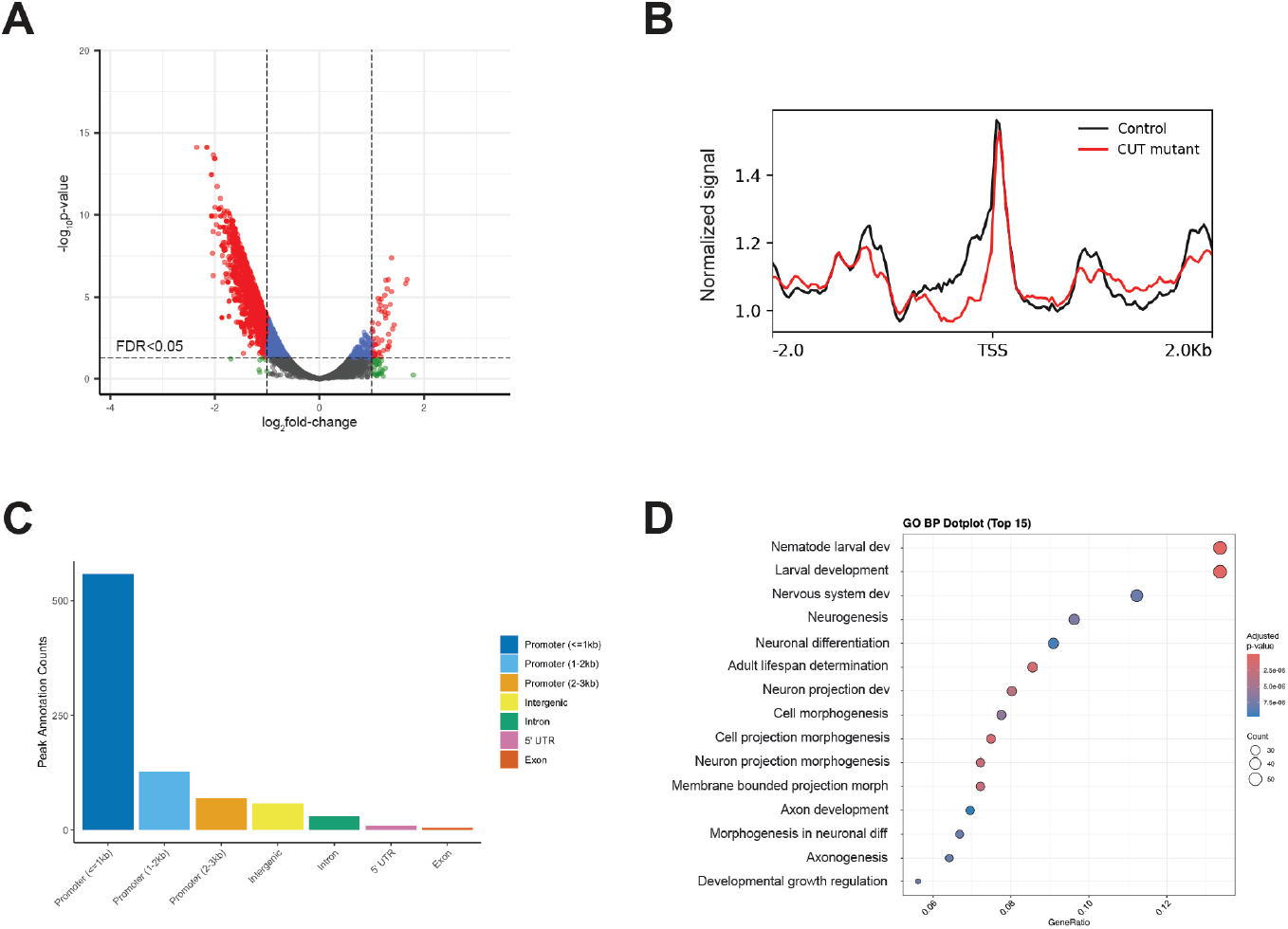
CUT-dependent chromatin accessibility changes in neuronal nuclei. **(A)** Volcano plot showing differential accessibility between Control and CUT mutant neuronal nuclei. Negative log2FC (left) indicates reduced accessibility in the CUT mutant, while positive log2FC (right) indicates increased accessibility in the CUT mutant (ATAC-seq, n = 2 biological replicates per genotype). Each point represents an ATAC peak, and points are colored by statistical and effect-size thresholds: differentially accessible peaks (FDR < 0.05 and |log2FC| > 1; red), significant but smaller effect (FDR < 0.05 and |log2FC| ≤ 1; blue), large effect without significance (FDR > 0.05 and |log2FC| > 1; green), and non-significant/small effect (grey). **(B)** Aggregate ATAC-seq signal centered on annotated TSSs for Control and CUT mutant samples. **(C)** Genomic feature annotation of differentially accessible peaks, including promoter-proximal bins defined as 0-1 kb, 1-2 kb, and 2-3 kb upstream of the TSS. **(D)** Gene Ontology (GO) biological process enrichment analysis for DA genes (n = 785 genes).

To determine where CUT-dependent accessibility changes occur, we annotated DA peaks by genomic feature. A substantial fraction of DA peaks mapped to promoter-proximal regions, with the remainder distributed across intronic and intergenic regions (**Fig. 2C**), indicating that CUT loss is associated with accessibility changes affecting promoter-proximal regulatory regions. To relate CUT-dependent accessibility changes to gene function, we defined CUT-dependent (DA) genes as unique genes assigned to DA peaks by the nearest annotated TSS within ±3 kb. Using this approach, we assigned the 858 DA peaks to 785 DA genes and performed gene ontology enrichment analysis on this gene set. This analysis highlighted enrichment for terms related to neuronal function and development (e.g., nervous system development, neuronal differentiation, and axon development), consistent with CUT-dependent accessibility changes impacting genes with neuronal roles (**Fig. 2D**). Together, these analyses indicate that CUT loss is associated with chromatin accessibility changes at a subset of promoter-proximal neuronal regulatory regions.

### Differentially accessible regions map to neuron-enriched and pan-neuronal genes

To contextualize CUT-dependent accessibility changes with respect to neuronal gene identity, we compared DA genes to reference neuronal gene categories. Specifically, we intersected DA genes with (i) neuronally enriched genes defined in our prior work (Leyva-Diaz and Hobert 2022) and (ii) a pan-neuronal gene set derived from CeNGEN single-cell expression data (Taylor et al. 2021) (**Fig. 3A**). This analysis revealed that a substantial fraction of DA genes overlap neuronally enriched and pan-neuronal gene sets, consistent with CUT-dependent accessibility changes affecting genes with neuronal expression programs.

**Figure 3.**
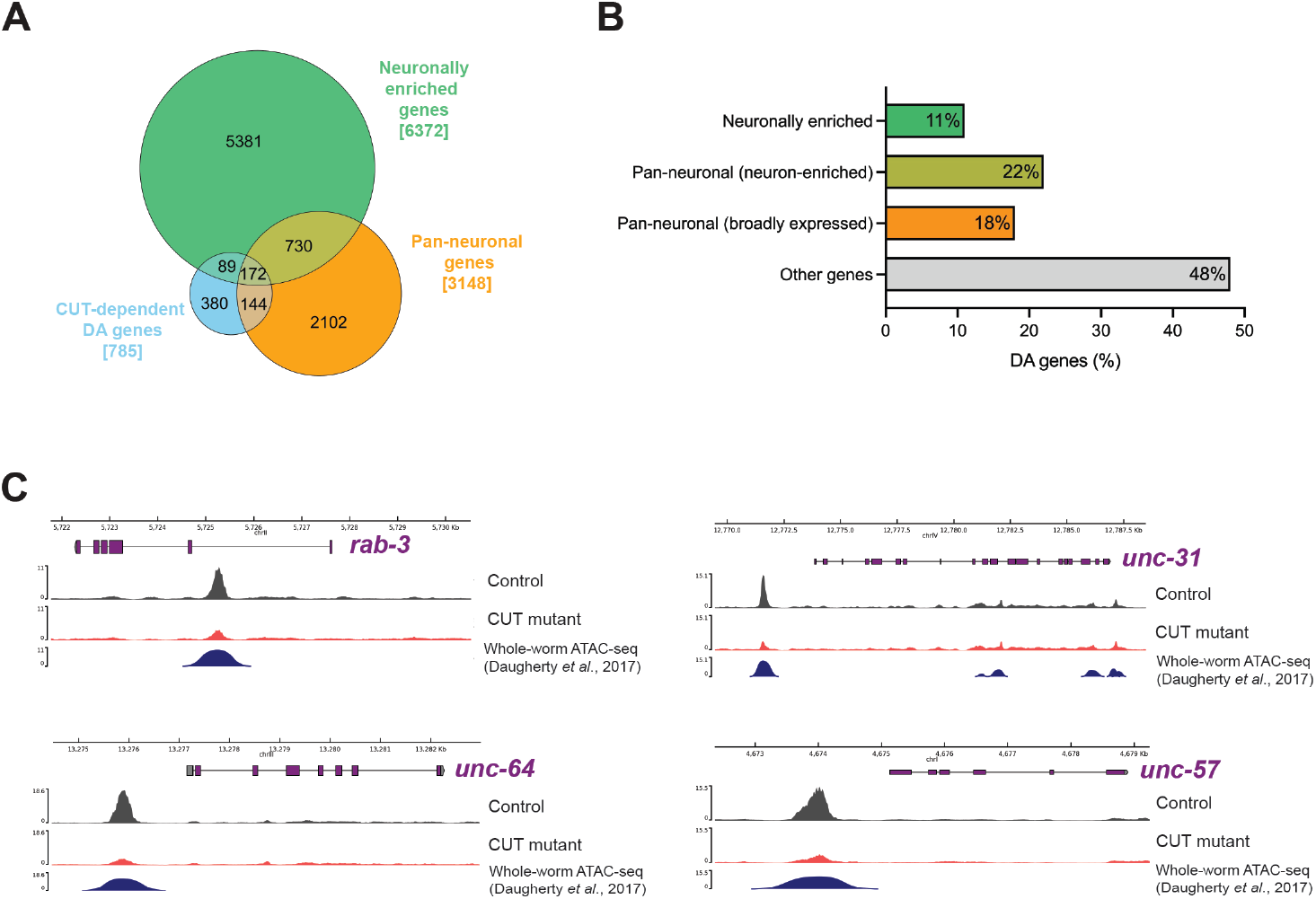
Differentially accessible peaks map to neuronally enriched and pan-neuronal genes. **(A)** Euler diagram showing overlaps between genes assigned to CUT-dependent differentially accessible ATAC-seq peaks (n = 785 genes), neuronally enriched genes (n = 6372 genes; Leyva-Diaz and Hobert 2022), and a curated pan-neuronal gene set derived from CeNGEN single-cell RNA-seq data (n = 3148 genes; Taylor et al. 2021). **(B)** Distribution of CUT-dependent DA genes across gene categories. Percentages indicate the fraction of all DA genes (n = 785) that overlap neuronally enriched genes, pan-neuronal (neuron-enriched) genes (DA genes overlapping the pan-neuronal set and also overlapping the neuronally enriched set), pan-neuronal (broadly expressed) genes (DA genes overlapping the pan-neuronal set but not the neuronally enriched set), or genes not assigned to the reference sets (“Other genes”). **(C)** Genome browser views of ATAC-seq signal at representative pan-neuronal loci (*rab-3/RAB3, unc-64/Syntaxin, unc-31/CADPS2, unc-57/SH3GL3*) showing Control and CUT mutant neuronal nuclei tracks (replicate averages) together with a published whole-worm reference ATAC-seq track (Daugherty et al. 2017; L3 stage).

To summarize these overlaps, we binned DA genes into four categories based on set membership (**Fig. 3B**). “Neuronally enriched” denotes DA genes overlapping the neuronally enriched reference set. “Pan-neuronal (neuron-enriched)” denotes DA genes overlapping the CeNGEN-derived pan-neuronal set and also classified as neuronally enriched (i.e., neuron-enriched pan-neuronal genes). A second subset of CeNGEN pan-neuronal genes did not overlap the neuronally enriched set and we refer to these as “Pan-neuronal (broadly expressed)”, in line with prior definitions distinguishing neuron-enriched versus broadly expressed pan-neuronal genes (Stefanakis et al. 2015). Because these categories are defined by overlap between reference gene sets, the “Pan-neuronal (neuron-enriched)” versus “Pan-neuronal (broadly expressed)” distinction is an operational classification rather than a direct measurement of tissue specificity. Remaining DA genes that did not fall into these categories were labeled “Other genes”. Within the DA gene set, 11% (n = 89 genes) were neuronally enriched, 22% (n = 172) were pan-neuronal (neuron-enriched), 18% (n = 144) were pan-neuronal (broadly expressed), and the remaining 48% (n = 380) were categorized as other genes (**Fig. 3B**). Notably, 52% of DA genes overlapped at least one neuronal reference category (neuronally enriched and/or pan-neuronal). Together, these annotations show that CUT-dependent accessibility changes include a substantial fraction of genes with established neuronal expression categories.

As an initial annotation step, we asked whether DA genes within the “neuronally enriched” and “pan-neuronal (neuron-enriched)” categories include genes with limited functional characterization. We operationally classified genes as “named” (gene name) versus “sequence-named” (systematic identifier) as a coarse proxy for annotation status. Within both subsets, a fraction of genes fell into the sequence-named category (**Fig. S1A**). To provide an expression-level context for these candidates, we examined their reported neuron-type expression patterns using the CeNGEN resource, visualizing expression for sequence-named candidates within the “neuronally enriched” subset (**Fig. S1B**) and the “pan-neuronal (neuron-enriched)” subset (**Fig. S1C**). These exploratory views are intended as descriptive context rather than functional inference. Together, these annotations provide a starting point to prioritize less-characterized candidates within neuronal expression categories for future functional analysis.

Finally, to illustrate CUT-dependent accessibility changes at representative neuronal loci, we visualized genome browser tracks at canonical pan-neuronal genes, including *rab-3/RAB3, unc-64/Syntaxin, unc-31/CADPS2*, and *unc-57/SH3GL3* (**Fig. 3C**). Gene views include a published whole-worm ATAC-seq reference track (Daugherty et al. 2017) to provide context for the accessible regions shown. At these loci, CUT mutant neuronal nuclei showed reduced accessibility relative to Control at promoter regions or nearby regulatory elements, consistent with the global bias toward reduced accessibility observed across DA sites (**Figs. 2A, 3C**). Together, these analyses connect CUT-dependent accessibility changes to neuronal gene programs and provide locus-level examples at established pan-neuronal genes.

## DISCUSSION

Our neuronal ATAC-seq profiling identifies widespread CUT-dependent changes in chromatin accessibility *in vivo*, with a strong bias toward decreased accessibility in CUT mutant neuronal nuclei. This directional shift is consistent with a contribution of CUT homeobox activity to supporting chromatin accessibility at a subset of neuronal regulatory regions, rather than causing symmetric gains and losses across the genome. CUT-dependent differentially accessible regions were frequently promoter-proximal and were associated with genes linked to neuronal expression categories, including neuronally enriched and pan-neuronal gene sets. This pattern suggests that CUT-dependent accessibility changes occur at regulatory regions positioned to impact gene regulation at neuronal genes. At the same time, a substantial fraction of DA genes fell outside current reference categories, highlighting both the breadth of CUT-dependent effects and the limited coverage of these reference gene sets.

Genome browser views at canonical pan-neuronal loci provide locus-level examples of CUT-dependent accessibility changes, with CUT mutant neuronal nuclei showing reduced accessibility relative to Control at promoters and nearby regulatory elements (**Fig. 3C**). We summarize these observations in a working model in which CUT activity helps support an accessible chromatin state at neuronal regulatory regions, thereby facilitating productive binding of additional transcription factors at shared targets (**Fig. 4**). In this view, loss of CUT function reduces accessibility at a subset of loci, which may in turn lower transcription factor occupancy and consequently reduce expression of CUT-dependent neuronal genes.

**Figure 4.**
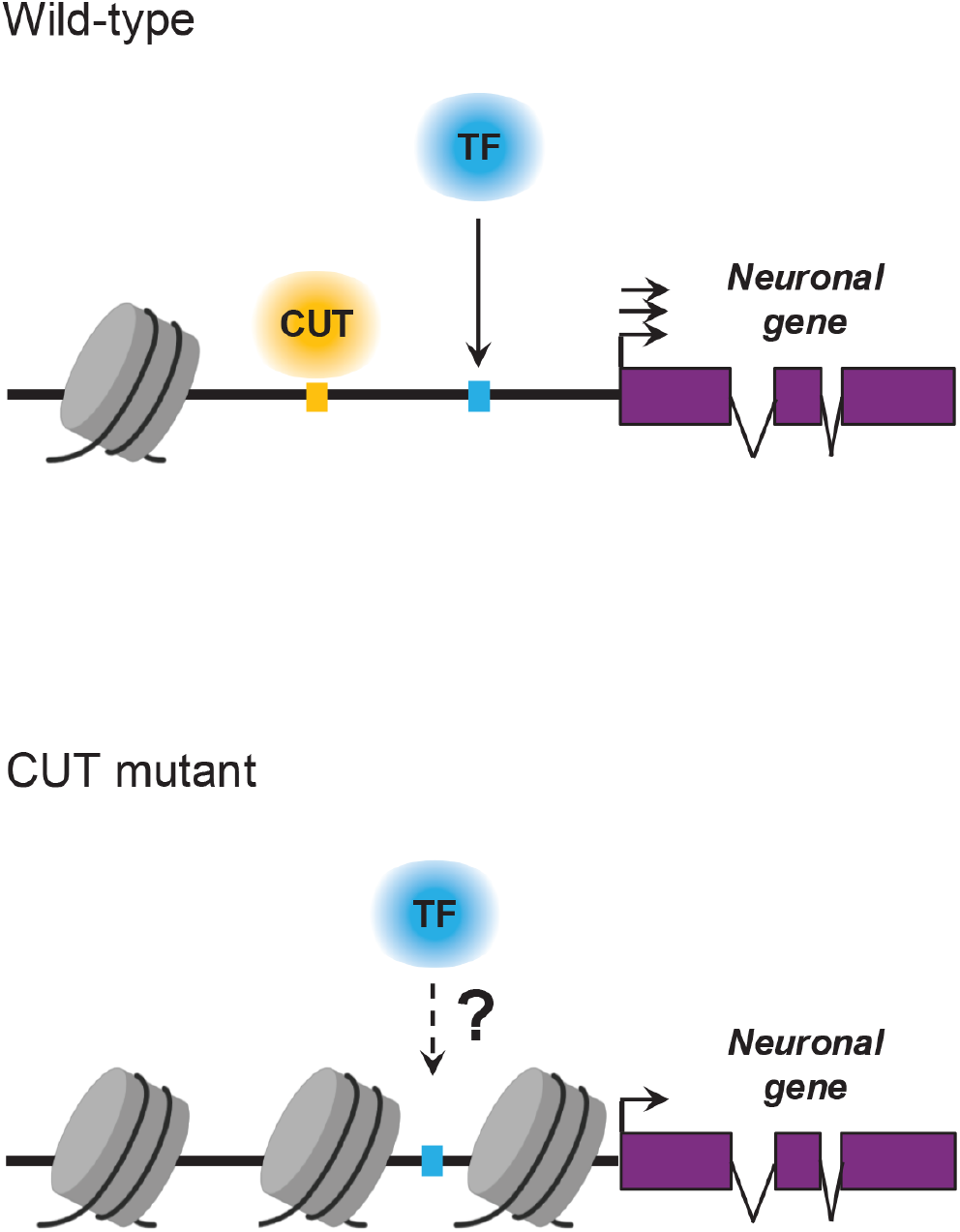
Working model for CUT-dependent neuronal chromatin accessibility. In wild-type neuronal nuclei, CUT homeobox factors support an accessible chromatin state at a subset of neuronal regulatory regions, facilitating occupancy by additional transcription factors. In CUT mutant nuclei, accessibility is reduced at these loci, which may reduce transcription factor occupancy and consequently lower gene output at CUT-dependent neuronal genes.

To place CUT-dependent accessibility changes in a broader regulatory context, we compared DA genes to independent datasets (**Fig. S2**). A majority of DA genes (518/785; 66%) overlap prior CUT binding evidence from ENCODE datasets (Luo et al. 2020; Kagda et al. 2025), consistent with CUT factors binding near many genes whose accessibility changes in the mutant. Because these comparisons are gene-level, they do not indicate that CUT binds the specific DA region assigned to each gene. We next compared DA genes to CUT-dependent transcriptional changes from our prior neuron-enriched bulk RNA-seq analysis in the same genetic backgrounds (Leyva-Diaz and Hobert 2022). By contrast, only a minority of DA genes overlap CUT-dependent differentially expressed genes in that dataset (100/785; 13%). This limited overlap suggests that CUT-dependent accessibility changes do not necessarily translate into detectable steady-state expression differences in bulk measurements and may reflect buffered, context-dependent, or neuron-subtype-restricted effects.

A limitation of bulk neuronal ATAC-seq is that it averages accessibility across all neuronal nuclei, leaving open whether CUT-dependent changes are broadly shared across the nervous system or concentrated in specific neuronal subtypes and regulatory programs. In addition, while our model proposes that CUT activity supports accessibility at neuronal regulatory regions in a manner that facilitates binding by other transcription factors (**Fig. 4**), directly evaluating this mechanism will require transcription factor-binding assays in neuronal nuclei and, ideally, neuron type-resolved measurements. More generally, resolving locus- and cell-type-specific effects and connecting accessibility changes to candidate partner factors will likely require single-nucleus chromatin profiling approaches. Together, the FANS-based neuron-enrichment strategy and datasets reported here provide a practical entry point for neuronal genomic profiling in *C. elegans* and a framework for interpreting CUT-dependent regulatory variation at the level of chromatin accessibility.

## MATERIALS AND METHODS

### Caenorhabditis elegans strains and handling

Worms were maintained under standard conditions at 20 °C on nematode growth medium (NGM) plates seeded with *E. coli* OP50. The wild-type strain used was Bristol variety, strain N2. The Control reporter strain was OH16748 (Sun and Hobert 2021), genotype *otIs790[UPN::npp-9::mCherry::blrp::3xFlag]*. The CUT mutant reporter strain was OH17055 (Leyva-Diaz and Hobert 2022), genotype *ceh-38(tm321) II; ceh-44(ot1028) III; ceh-48(tm6112) IV; otDf1 X; otIs790*. The *otIs790* transgene expresses a pan-neuronal INTACT-style nuclear envelope tag under the control of the synthetic ultra pan-neuronal (UPN) promoter (Yemini et al. 2021), driving expression of the NPP-9 nucleoporin fused to mCherry, and to BLRP/3xFLAG for affinity purification protocols (Steiner et al. 2012; Sun and Hobert 2021).

### Neuronal nuclei isolation and FANS sorting

Synchronized L1 larvae were obtained by standard bleach-based egg preparation. Worms were washed to remove bacteria and resuspended in ice-cold nuclei isolation buffer (10 mM Tris-HCl pH 7.5, 10 mM NaCl, 10 mM KCl, 2 mM EDTA, 0.5 mM EGTA, 0.5 mM spermidine, 0.2 mM spermine, 0.2 mM DTT, 0.1% NP-40). Nuclei were released using a bead-assisted lysis workflow combined with freeze-thaw, detergent-based permeabilization, and glass dounce homogenization. Lysates were cleared by low-speed centrifugation to remove worm debris (100g, 1 min, 4°C), and nuclei were stained with DAPI prior to sorting. Samples were kept on ice throughout handling and transport. Nuclei were sorted on a BD FACS Discover S8 instrument by FANS using an 86 µm nozzle. Gating was performed sequentially to exclude debris based on scatter parameters, select DAPI(+) events, and collect the mCherry high population corresponding to neuronal nuclei. The mCherry positivity gate was defined using N2 wild-type animals lacking the reporter. For ATAC-seq, nuclei from ∼50,000 L1 animals were sorted into a single tube (BSA-coated), which was then centrifuged to concentrate nuclei for downstream processing (500 g, 5 min, 4°C). Nuclei integrity was monitored by fluorescence microscopy of DAPI-stained aliquots before sorting, immediately after sorting, and after post-sort centrifugation using a Leica DM5000B microscope. Images were processed in ImageJ software.

### ATAC-seq library preparation and sequencing

ATAC-seq libraries were prepared from FANS-sorted mCherry(+) neuronal nuclei (∼100,000 nuclei per library). Tagmentation was performed using Diagenode loaded Tn5 transposase. Libraries were PCR-amplified using NEBNext Ultra II Q5 Master Mix (New England Biolabs), with the number of cycles determined by qPCR-based cycle estimation using SYBR Fast 2x PCR Master Mix (ABI Prism). Libraries were purified and size-selected using AMPure XP beads (Beckman Coulter) and assessed by Qubit quantification and Bioanalyzer fragment analysis. Libraries were sequenced by Novogene on an Illumina NovaSeq X Plus platform (paired-end 150 bp).

### ATAC-seq data analysis

ATAC-seq computational analyses were performed in R (v4.5.0). Paired-end BAM files from Control replicates and CUT mutant replicates were quantified over a consensus peak set by converting the BED peak coordinates to SAF and counting paired-end fragments (read pairs) per peak with Rsubread featureCounts (v2.22.1). A consensus peak atlas was generated as the union of per-sample Genrich peak calls, with overlapping intervals merged using the IRanges package to produce a non-redundant reference set. Differential accessibility was tested with DESeq2 (v1.48.1) using a negative binomial model (Control as reference), after filtering out low-count peaks (total counts ≤10 across samples). Differentially accessible peaks were defined as those with FDR (BH-adjusted p-value) < 0.05 and |log2FC| > 1. DA peaks were annotated with ChIPseeker (v1.44.0) using a transcript database built from the *C. elegans* ce10 (WS220) GTF via GenomicFeatures (v1.60.0), defining promoter-proximal regions as ±3 kb around the TSS. For gene-level analyses, peaks were assigned to the nearest annotated TSS within ±3 kb. Genome-wide coverage tracks were generated as bigWig files from BAM files using deepTools bamCoverage (v3.5.1). Track visualizations were generated with pyGenomeTracks using the corresponding bigWig files. Gene Ontology enrichment (Biological Process) on DA genes was performed with clusterProfiler (v4.16.0) and org.Ce.eg.db (v3.21.0), with visualization using enrichplot (v1.28.2). Plots were generated with ggplot2 (v3.5.2) and volcano plots with EnhancedVolcano (v1.26.0).

### RNA extraction and RT-qPCR

Total RNA was extracted from sorted neuronal nuclei fractions (mCherry(+) and mCherry(-)) using the NucleoSpin RNA XS kit (Macherey-Nagel) according to the manufacturer’s instructions, including on-column DNase digestion. cDNA was synthesized using the SuperScript IV First-Strand Synthesis System (Thermo Fisher Scientific) with a 1:1 mixture of oligo(dT) and random hexamer primers. RT-qPCR was performed using SYBR Green chemistry (SYBR Fast 2x PCR Master Mix; KAPA Biosystems) on a StepOne Real-Time PCR System (Applied Biosystems). Expression levels of *egl-3* and *myo-3* were quantified and normalized to *Y45F10D*.*4* using the ΔΔCt method, with the mCherry(-) fraction used as the calibrator (set to 1). Biological replicates corresponded to independent sorts (n = 3), with two technical qPCR replicates per sample. Statistical significance was assessed in GraphPad Prism using an unpaired two-tailed t test on biological replicate values.

### Gene set definitions and sources

Neuronally enriched genes were taken directly from the published gene lists in Leyva-Diaz and Hobert 2022, where enrichment was defined by DESeq2 comparing wild-type neuronal nuclei immunoprecipitated (IP) samples versus total nuclei input (FDR < 0.05) and neuronally enriched genes were those with log2FoldChange > 0.

CUT-dependent differentially expressed (DE) genes were taken directly from the published differential expression results in Leyva-Diaz and Hobert 2022 (DESeq2; FDR < 0.05; all DE genes, including up- and downregulated genes).

CUT target genes were taken from previously published CEH-48 and CEH-38 ChIP-seq datasets (CEH-48: ENCODE experiment ENCSR844VCY; CEH-38: modENCODE accession modEncode_4800), and we used the target-gene lists as reported in Leyva-Diaz and Hobert 2022. In that study, ChIP peaks were intersected with promoter regions (5 kb upstream to 1 kb downstream of the annotated TSS), and overlapping genes were called as targets. The union of CEH-48 and CEH-38 target genes was used for overlap analyses.

A curated pan-neuronal gene set was derived from CeNGEN scRNA-seq neuron-class expression data (Taylor et al. 2021). A strict requirement for detectable expression in 100% of neuron classes yielded only 207 genes, consistent with under-detection of broadly but lowly expressed transcripts due to dropout and limited sequencing depth. To calibrate an operational “pan-neuronal” threshold, we used a core set of 23 validated pan-neuronal genes defined by fosmid reporter expression (Stefanakis et al. 2015) and asked under which conditions these genes are consistently detected in CeNGEN. We selected neuron classes represented by ≥99 cells (103 neuron classes) as a high-depth subset and defined pan-neuronal candidates as genes with detectable expression in ≥97/103 neuron classes (94%). This empirically chosen compromise recovers the validated pan-neuronal reporters while limiting inclusion of ubiquitously expressed genes. The resulting curated pan-neuronal gene set contained 3,148 genes and was used for downstream analyses.

All reference gene sets used in this study (neuronally enriched genes, CUT-dependent RNA-seq DE genes, CUT ChIP-seq target genes, and the curated pan-neuronal gene set) are provided in **Table S1**. Gene identifiers were harmonized using WormMine and all set intersections were performed using WormBase gene IDs (WS298).

### Gene nomenclature classification and CeNGEN expression visualization

“Uncharacterized” genes were operationally defined as genes lacking a standard gene name and annotated only by a systematic sequence identifier (sequence-named genes). Gene names were harmonized using WormMine and analyses were performed using WormBase gene IDs (WS298). Uncharacterized genes were identified within CUT-dependent DA genes intersecting the neuronally enriched set (“neuronally enriched” in **Fig. 3B**) and within the intersection of CUT-dependent DA genes with both the neuronally enriched and curated pan-neuronal sets (“pan-neuronal (neuron-enriched)” in **Fig. 3B**). CeNGEN expression dot plots were generated using the CengenApp web application (Taylor et al. 2021; https://cengen.shinyapps.io/CengenApp/), using the “Heatmaps of gene expression” option and the neuron-class dataset “Neurons (threshold 2)” (full neuron-class list). Dot color (scaled TPM) and dot size (fraction of cells with detectable expression) are displayed as defined by CeNGEN. Dot sizes are scaled within each panel, as defined by CeNGEN, and are not quantitatively comparable between panels.

### Use of generative AI tools

A generative AI tool was used to assist with language editing and improving clarity of the manuscript. All content was reviewed and approved by the authors, who take full responsibility for the accuracy, integrity, and originality of the work.

## ACKNOWLEDGEMENTS

We thank members of the López-Bendito lab and Flames lab for comments, Ser van der Burght (Ahringer lab) for ATAC-seq technical advice, WormBase (Sternberg et al. 2024) and the CGC (funded by the NIH Office of Research Infrastructure Programs, P40 OD010440) for providing resources and reagents. Some figure elements were created with BioRender.com. This work was funded by a CIDEGENT grant from Generalitat Valenciana (CIDEXG/2022/30) (E.L.-D.).

## AUTHOR CONTRIBUTIONS

E.L.-D. conceived the project, designed the experiments and prepared the manuscript. J.I.J.-L., J.V., and E.L.-D. performed the experiments and analyses.

## FIGURES

**Supplementary Figure S1.**
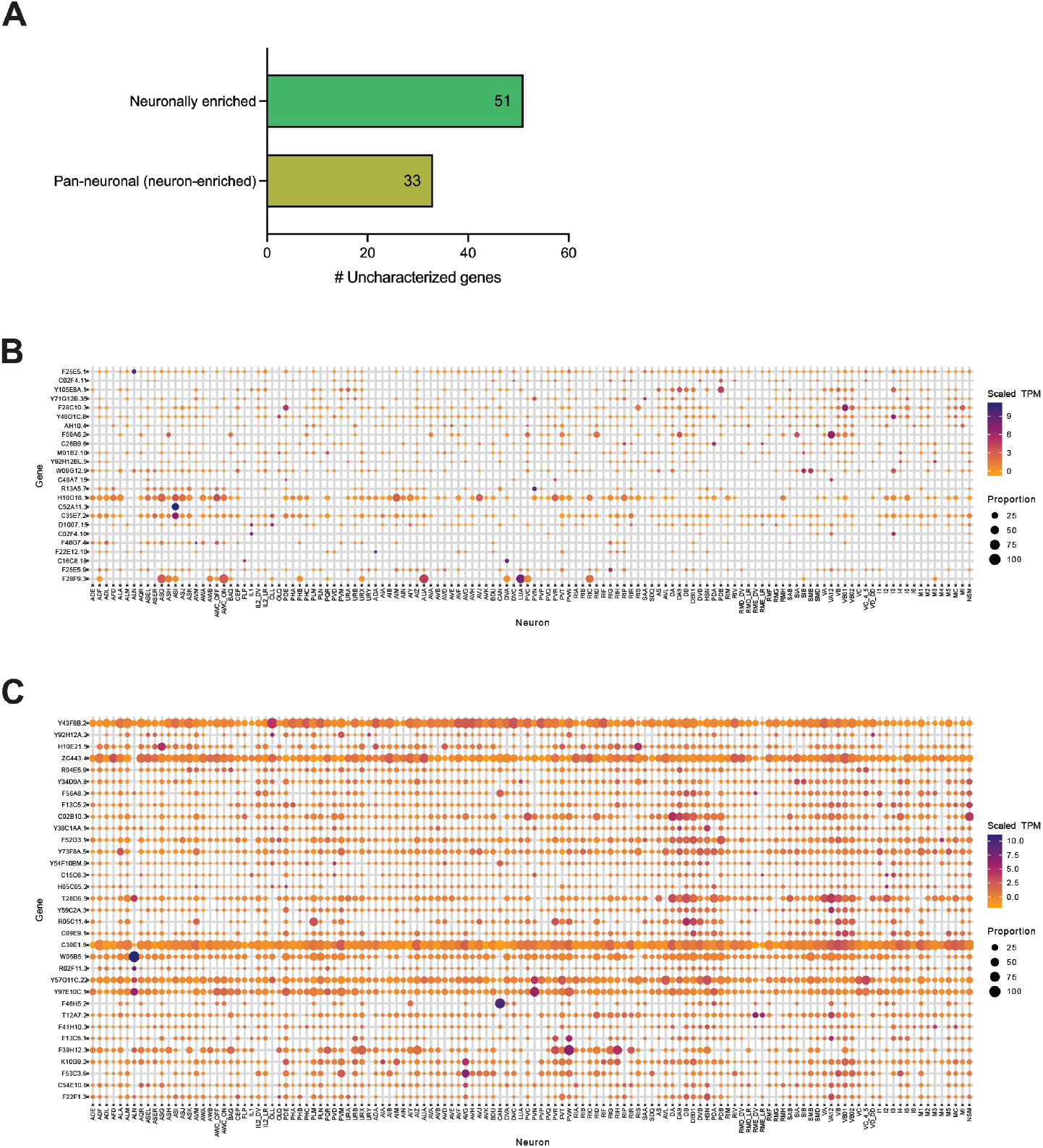
Uncharacterized (sequence-named) DA genes and CeNGEN expression context. **(A)** DA genes were classified as uncharacterized (systematic locus identifier lacking a standard gene symbol) within the “neuronally enriched” and “pan-neuronal (neuron-enriched)” subsets defined in **Fig. 3B**. **(B-C)** CeNGEN expression dot plots for uncharacterized DA genes in the “neuronally enriched” subset (**B**; 24 genes shown) and the “pan-neuronal (neuron-enriched)”subset (**C**; 33 genes shown). Dot color indicates scaled TPM and dot size indicates the fraction of cells with detectable expression, as defined by CeNGEN (Taylor et al. 2021). Dot sizes are scaled within each panel and should not be compared quantitatively between panels. In **(B)**, 27 genes from the “neuronally enriched” subset are not shown because no signal is displayed under these CeNGEN settings.

**Supplementary Figure S2.**
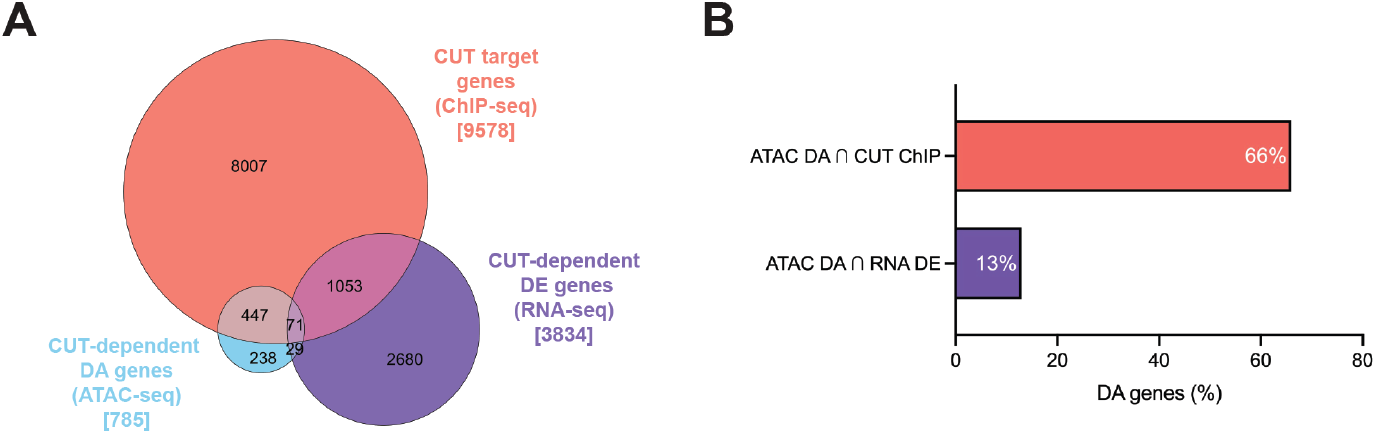
Comparison of CUT-dependent ATAC-seq changes with prior CUT binding and neuronal RNA-seq. **(A)** Euler diagram showing overlaps among CUT-dependent DA genes (ATAC-seq; n= 785), CUT-dependent DE genes (RNA-seq; n = 3834; Leyva-Diaz and Hobert 2022), and CUT target genes (ChIP-seq; n = 9578; union of CEH-48 and CEH-38 target gene sets extracted from ENCODE datasets (Luo et al. 2020; Kagda et al. 2025)). **(B)** Percentage of CUT-dependent DA genes overlapping the CUT target gene set (ChIP-seq; 66%, 518 genes) or the CUT-dependent DE gene set (RNA-seq; 13%, 100 genes).

